# Differential Dynamics and Roles of FKBP51 Isoforms and Their Implications for Targeted Therapies

**DOI:** 10.1101/2024.08.03.606475

**Authors:** Silvia Martinelli, Kathrin Hafner, Maik Koedel, Janine Knauer-Arloth, Nils C Gassen, Elisabeth B Binder

## Abstract

The expression of FKBP5, and its resulting protein FKBP51, is strongly induced by stress and glucocorticoids. Numerous studies have explored their involvement in a plethora of cellular processes and diseases, including psychiatric disorders, inflammatory conditions and cancer. However, there is a lack of knowledge on the role of the different RNA splicing variants and the two protein isoforms that originate from the human FKBP5 locus, especially in response to glucocorticoids. In this study we use *in vitro* models as well as peripheral blood cells of a human cohort to show that the two expressed variants are both dynamically upregulated following dexamethasone. We also investigate the subcellular localization of the protein isoforms, their degradation dynamics as well as their differential role in known cellular pathways. The results shed light on the difference of the two variants and highlight the importance of differential analyses in future studies with implications for targeted drug design.

## Introduction

The FK506 binding protein 51 (FKBP51) is a ubiquitously expressed immunophilin, encoded by the gene *FKBP5*, and whose function has been investigated in association with numerous biological processes describing FKBP51 as a central regulator of pathways involved in psychiatric and neurodegenerative disorders, immune response, inflammation, cardiovascular diseases, metabolic pathways and cancer (Zannas *et al*, 2016; Blair *et al*, 2015; Marrone *et al*, 2023b; Smedlund *et al*, 2021; Zannas *et al*, 2019). The most investigated role of FKBP51, is its involvement in the regulation of the stress response, initially discovered in squirrel monkeys where it was observed that an increased expression of FKBP51 is the cause of glucocorticoid receptor (GR) resistance and high circulating cortisol levels in these animals (Denny *et al*, 2000; Scammell *et al*, 2001). The critical role of FKBP5 in hypothalamus-pituitary-adrenal (HPA) axis regulation is also supported by the fact that feedback control of this axis is impaired in FKBP51-deficient mice. In fact, FKBP51 is an inhibitor of the GR, the key effector of the hypothalamus-pituitary-adrenal (HPA) axis. On the other hand, it is also a transcription target of the GR, with several glucocorticoid response elements (GREs) in intronic and upstream enhancer regions and strong upregulation observed across many tissues. This can lead to an ultra-short negative feedback of GR activity (Jääskeläinen *et al*, 2011). FKBP51 is a co-chaperone protein and interacts not only with the GR via HSP90, but also with many other proteins, via direct protein-protein interactions (Taipale *et al*, 2014; Martinelli *et al*, 2021). These interaction partners include heat shock proteins, steroids receptors, PH domain and leucine-rich repeat protein phosphatases (PHLPP) and Akt, Nuclear Factor ‘Kappa-Light-Chain-Enhancer’ of Activated B-Cells (NF-kB) as well as DNA methyltransferase 1, Calcineurin-NFAT signaling, Tau and others (Hähle *et al*, 2019). These interactions have been shown to play a relevant role in many cell types, including cancer cells (Pei *et al*, 2009; Hähle *et al*, 2019). Given the strong upregulation of FKBP51 following glucocorticoid exposure and its many downstream partners and thus central role promoting a cellular stress response, it is a particularly interesting potential drug target for a number of diseases.

In humans, four transcription variants (variant 1-4) of *FKBP5* have been annotated, coding for two different isoforms (isoform 1 and 2) of the protein FKBP51. In mice, a species from which a large body of current knowledge on FKBP5/51 is derived, only one isoform (corresponding to the human isoform 1) of FKBP51 is annotated. The human transcription variant 1 (ENST00000357266.8) differs in the 5 ’UTR from variants 2 (ENST00000536438.5) and 3 (ENST00000539068.5) and all three code for the 475 amino acid (aa) long isoform 1 (Q13451-1), while variant 4 (ENST00000542713.1) corresponds to a much shorter transcript and codes for the “truncated” isoform 2 (Q13451-2) of 268 aa (see Fig. 1 a and b). Gene expression data (Lonsdale et al, 2013, www.gtexportal.org) indicate that *FKBP5* is ubiquitously expressed with particularly high expression levels in tibial nerve, skeletal muscle and esophagus (Fig. 1 c). Variant 1 is more strongly expressed than the other variants, suggesting that in all experiments conducted without distinguishing between the different variants, the overall expression levels mirror mainly the ones of variant 1. Variant 2 appears to be the least expressed, while it doesn’t seem to be a quantitative difference between variant 3 and 4, but rather a tissue-specific differential expression.

**Figure 1.**
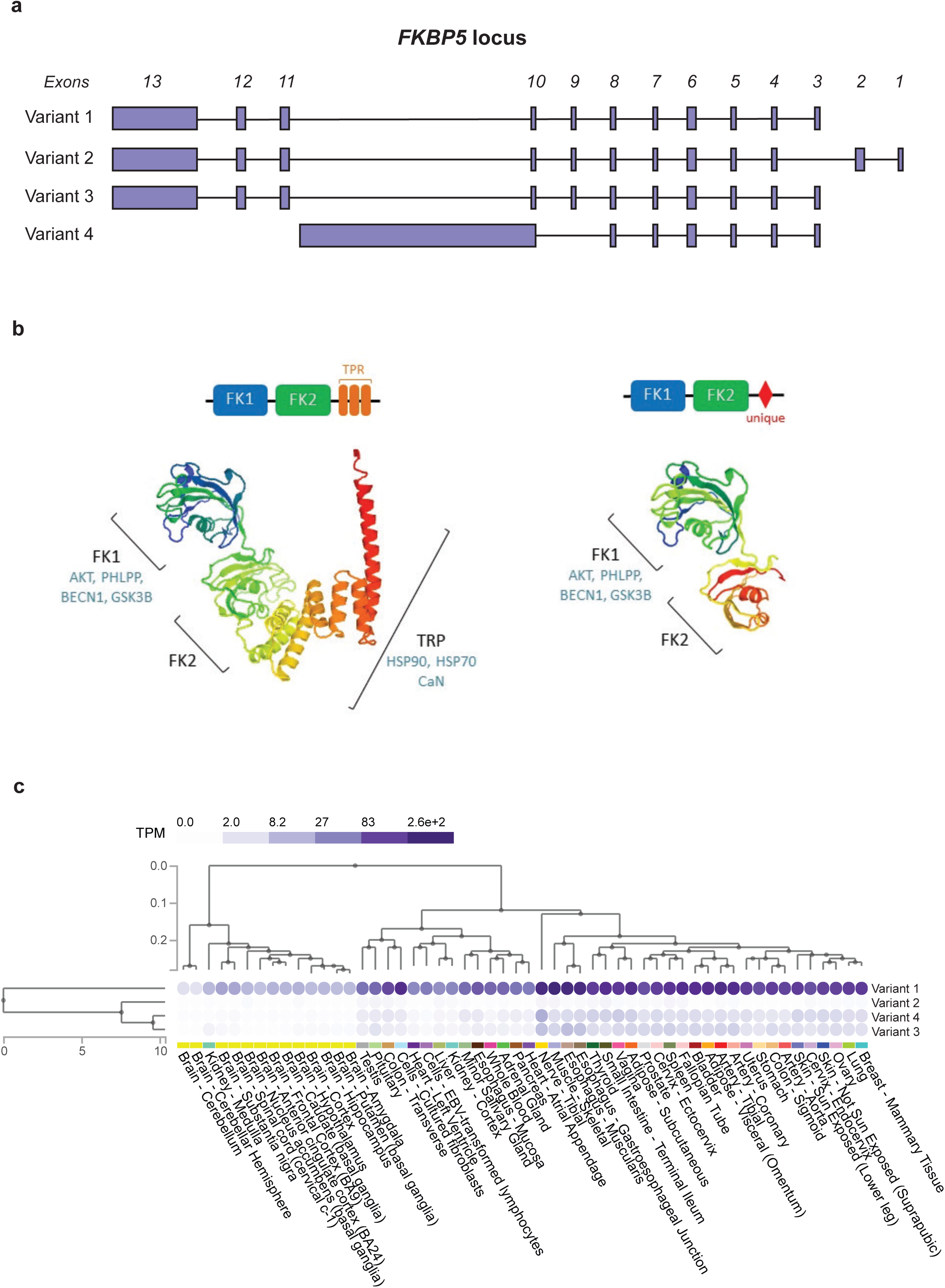
FKBP5/51 transcription variants and isoforms. a) Schematic view of the FKBP5 locus on human chromosome 6 and the four splicing variants of the gene (adapted from gtexportal.org). b) Schematic view of FKBP51 isoform 1 and 2 protein structures and 3D structure models generated with the swiss model repository server of the expasy portal (https://swissmodel.expasy.org/). Domains are indicated in black and experimentally validated domain-associated binding partners in blue. c) Transcription variant-specific FKBP5 expression throughout human tissues (adapted from gtexportal.org).

At the protein level, FKBP51 isoform 1 has two N-terminal FK506-binding (FK) domains and three tetratricopeptide repeat (TPR) motifs at the C-terminus (Fig. 1 b). Isoform 2 of FKBP51 shows sequence identity with isoform 1 for the first 222 aa, corresponding to the FK domains. The sequence ranging from aa 223 to its C-terminus (aa 268) is unique and, so far, uncharacterized. Missing the rest of isoform 1’s C-terminal region, isoform 2, therefore, lacks the TPR motifs (Fig. 1 b). The first FK domain, FK 1, is the binding site of the immunosuppressive drug FK506, from which the protein gets its name. FK1 also exerts a peptidyl-prolyl cis-trans isomerase (PPIase) or rotamase activity (Schiene & Fischer, 2000), characteristic of all immunophilins. The pocket in FK1 is also the binding site for another drug, rapamycin. This drug, in complex with FKBP51, exerts immunosuppressive and anticancer effects, mediated via the selective inhibition of the mechanistic target of rapamycin or mTOR (Sabatini, 2006). Downstream, adjacent to FK1, is the second FK domain, FK2, that is presumably derived from a duplication event of the FK1 domain and shares 32% sequence homology with it (Cioffi *et al*, 2011), but lacks measurable rotamase activity (Sinars *et al*, 2003) and does not bind FK506. Instead, it might have cooperative functions with the TPR motifs (Sinars *et al*, 2003). The TPR motifs at the C-terminus promote protein-protein interactions (Russell *et al*, 1999), in particular with chaperone proteins such as HSP90 and heat shock protein 70 (HSP70) (Dornan *et al*, 2003). Furthermore, Li and colleagues showed that the TPR motifs are also responsible for the interaction with the serine-threonine phosphatase calcineurin (CaN) (Li *et al*, 2002). This phosphatase activates nuclear transcription factors of activated T lymphocytes (NFAT), responsible for the expression of interleukin-2 (IL2) and several T cell specific activators, regulating thereby the clonal expansion of T cells after stimulation by an antigen (Li *et al*, 2002). Thus far, only a few studies carried out by the group of M.F. Romano at the University of Naples, Italy, have described different functions of the human isoforms, with isoform 2 associated to the development of melanoma and glioma ((Romano *et al*, 2015; D’Arrigo *et al*, 2017)). Given the substantial structural difference of the two FKBP51 isoforms and the scarcity of studies regarding their differential roles, we decided to investigate possible differential functions of FKBP51 isoforms. We first mapped transcript and isoform differences in the expression dynamics following induction by glucocorticoids and then probed functional differences in key pathways. A better understanding of the function of different isoforms may help improving the development of FKBP51-targeting drugs.

## Results

### Expression and degradation dynamics of *FKBP5* / FKBP51

In order to characterize the expression dynamics of *FKBP5*, we first determined the expression via reverse transcription quantitative polymerase chain reaction (RT-qPCR) in HeLa cells. Results evidenced the absence of significant expression of variants 2 and 3. The probes covering all FKBP5 variants yielded the strongest signal, while variant 4 showed lower yet measurable levels (Fig. 2a, S1). This indicates that large part of this signal derives from variant 1 expression. Given the key role of *FKBP5* in the stress response, we were interested in the differential expression dynamics of the transcription variants upon GR activation. For this purpose, we stimulated HeLa cells with 100 nM of the GR agonist dexamethasone (Dex) for 2, 4, 6, 12 and 24 hours. Transcription levels of the different mRNA variants were subsequently analyzed via RT-qPCR (Fig. 2 b). Due to the lack of sequence uniqueness for variant 1, probes spanning variants 1, 2 and 3 were used. Considering the absence of variants 2 and 3, the observed signal was assumed to correspond to variant 1 and will be referred to as variant 1 from here on. The expression of both variant 1 and 4 was significantly increased in response to Dex across time (Two-way ANOVA, time effect p < 0.0001) and a significant difference between variants over time (two-way ANOVA, time x variant effect p < 0.0005). Interestingly, despite having lower expression at basal levels, variant 4 showed an increased response ratio over vehicle compared to variant 1 at early time points (significant difference at 2 and 4 hours). Furthermore variant 4 showed a more rapid response to Dex than variant 1: variant 4 levels were significantly increased already after two hours of treatment while at the same time point variant 1 was still expressed at baseline levels. After 6 hours treatment and until the end of the treatment period at 24 hours, both variants showed a significantly increased expression compared to baseline but with no difference between each other: variant 4 follows a steady slope after 6 hours treatment, while variant 1 expression reflects in a slowly increasing curve up to 24 hours.

**Figure 2.**
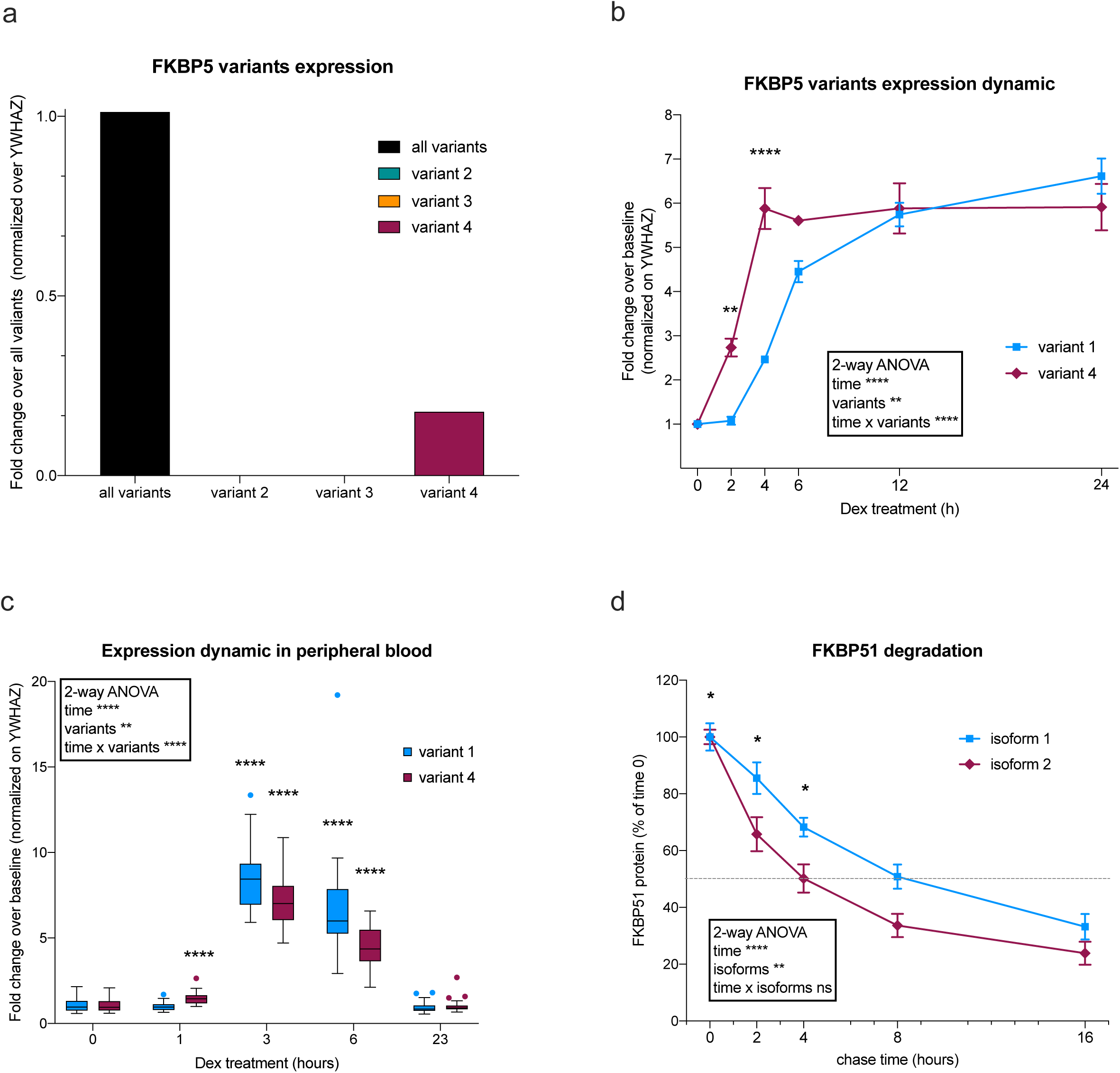
Expression of FKBP5 splicing variants in HeLa cells. a) RT-qPCR quantification of FKBP51 variants in unstimulated HeLa cells b) RT-qPCR quantification of FKBP51 variants, expressed as fold change of Dex-treated over vehicle-treated, normalized on the housekeeper YWHAZ, of HeLa cells treated with 100 nM Dex or vehicle for 24 hours. Two-way ANOVA with Geisser-Greenhouse correction (shown in the box) and Sidak’s multiple comparisons test (shown in the graph). Data shown as mean ± s.e.m. c) Fold change of FKBP5 Variants 1 and 4 over vehicle and normalized over YWHAZ at 0, 1, 3, 6 and 23 hours after Dex stimulation. Mixed effects model with Geisser-Greenhouse correction (shown in the box) and Sidak’s multiple comparisons test (shown in the graph). Data shown as box-and-whisker plot (Tukey style). d) Pulse chase assay of FKBP51 isoform 1 and 2 of HeLa cells transfected with HaloTag®-tagged-isoform 1 or HaloTag®-tagged-isoform 2, pulsed with a fluorophore and chased for 2, 4, 8, and 16 hours. Quantifications were made from western blots. *P < 0.05. Two-way ANOVA (shown in the box) and Sidak’s multiple comparisons test (shown in the graph). Data shown as mean ± s.e.m. For all statistics *P < 0.05, **P < 0.01, ****P < 0.0001.

Having seen a difference in the response to Dex between variant 1 and variant 4 in cell culture, we decided to assess whether this finding holds true in a different tissue *in viv*o. For this purpose, we analyzed via RT-qPCR the expression of FKBP5 variant 1 and variant 4 in peripheral blood samples of a cohort of 26 healthy, male participants at baseline and 1, 3, 6 and 23 hours after 1.5 mg Dex intake. The results showed a much lower expression of variant 4 compared to variant 1 as can be appreciated from the Ct values (Fig. S3), which is in accordance to data for peripheral blood from GTEX (gtexportal.org). Upon Dex stimulation, we observed an increased expression of both variants, but with a significantly different expression dynamic over the time course (Fig. 2c; two-way ANOVA time effect p < 0.0001 and time x variants effect p < 0.0001). While a significant increase was detected for both variants after 3 and 6 hours of Dex induction compared to baseline, variant 4 showed an increased expression already after one hour in contrast to variant 1, confirming the faster dynamic observed *in vitro* (Fig. 2c; two-way ANOVA variants effect p < 0.0058). Interestingly, in blood samples we can observe a faster dynamic compared to cell culture material, with a return to baseline levels after 23 hours.

After having observed a different expression dynamic of variant 1 and 4 over time in response to Dex, we investigated the half-life of the respective protein isoforms: isoform 1 and 2. For this purpose, a pulse-chase approach was used. HeLa cells were transfected with HaloTag®-tagged plasmids coding for either isoform 1 or 2. Twenty-four hours later, cells were tagged with a cell permeable halogenated fluorophore 16, 8, 4 and 2 hours before harvesting. After harvesting the cells, proteins were extracted and subjected to western blot, and fluorescence intensity was measured on nitrocellulose membrane. Results indicated that both isoforms are degraded throughout the 24 hours (two-way ANOVA time effect p = 0.0001) at a significantly different rate (two-way ANOVA isoform effect p < 0.0001 and time x isoform effect p < 0.0001). The degradation of isoform 2 is faster with a half-live of four hours, while isoform 1 reached 50% of degradation only after 8 hours (Fig. 2d). These results suggest a faster turnover of variant 4/isoform 2 with an increased and faster responsiveness to Dex and a shorter half-life of the protein compared to variant 1/ isoform 1.

### Differential regulation of cellular pathways

#### Subcellular location of the two isoforms

To better understand possible differences in their functions, we analyzed the intracellular localization of the two isoforms. Overall, the information about FKBP51’s intracellular localization appears to be highly dependent on antibodies used for detection, and no information is available for the different isoforms. To avoid potential artefacts deriving from immunocytochemical processing, HeLa cells were transfected with plasmids coding for GFP-tagged isoforms 1 or 2. A plasmid expressing only GFP was used as control. Cells were live imaged 24 hours after transfection. Resulting images (Fig. 3 a) showed ubiquitous signal from the control-transfected cells. Isoform 1 presented a cytoplasmic accumulation, while isoform 2 showed a distinct subnuclear localization. In support of this result, a motif analysis performed with the Expasy Prosite database (https://prosite.expasy.org/) revealed a possible bipartite nuclear localization signal (NLS) between aa 232 and 246 (supplementary Fig. S4), which corresponds to the region of the protein that is unique for isoforms 2 compared to isoform 1. In fact, despite having a low confidence level (score 3.000), the same analysis performed on isoform 1 could not detect any NLS (supplementary Fig. S2). Given the structural and sub cellular localization differences, we investigated whether these have a functional effect. To this aim, we investigated different cellular pathways that are known to be regulated by FKBP51, and analyzed the differential role of the two isoforms on them.

**Figure 3.**
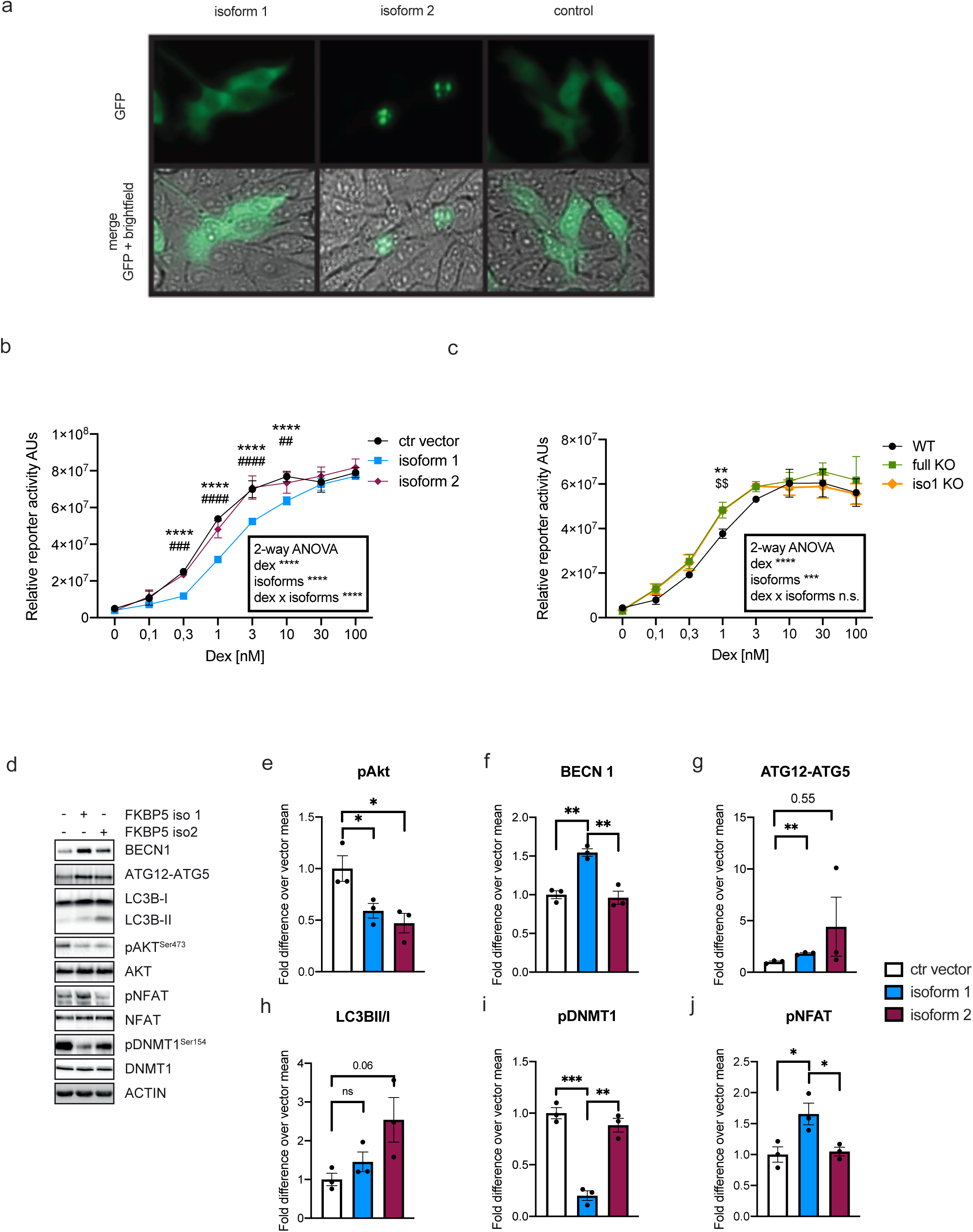
Differential pathway regulation of FKBP51 isoforms. **a)** Epifluorescent and bright field imaging of HeLa cells transfected with GFP-tagged FKBP51 isoform 1, GFP-tagged FKBP51 isoform 2, or GFP-control vector 24 hours prior to imaging. **b-c)** GRE-driven reporter gene assay performed in HeLa cells transfected with **b)** FKBP51 isoform 1, FKBP51 isoform 2 or an empty vector (ctr vector), or **c)** in WT, full KO and Isoform 1 KO (iso1 KO) HeLa cells treated with 0.1 nM, 0.3 nM, 1 nM, 3 nM, 10 nM, 30 nM, 100 nM or vehicle for 4 hours. Two-way ANOVA (shown in the box) with Tukey multiple comparisons test (shown in the graph). * indicates comparison with control/WT and isoform 1/full KO, # indicates comparison between isoform 1 and isoform 2, and $ refers to comparison between WT and iso 1 KO. *P < 0.05, **P < 0.01, ***P < 0.0005, ****P < 0.0001. **d-h)** Quantification of western blots analyses for different pathway markers from HeLa cells transfected with FKBP51 isoform 1, FKBP51 isoform 2 or an empty vector: **d)** phosphorylated AKT (pAKT) normalised on total AKT,**e-g)** autophagy markers, BECN1, ATG12 and LC3BII/I; **h)** phosphorylated DNMT (pDNMT) normalised on total DNMT; **i)** Quantification of western blots analyses for phosphorilated NFAT (pNFAT) normalised on total NFAT from Jurkat cells transfected with FKBP51 isoform 1, FKBP51 isoform 2 or an empty vector; *P < 0.05. Mann-Whitney test. * and # indicate comparisons with ctr vector and isoform 2 respectively. Data shown as mean ± s.e.m.

#### Differential effects on GR activity

As one of the best-known functions, we first analyzed the negative regulation of the two isoforms on GR. Activity of the different isoforms on GR was assessed via Glucocorticoid Response Element (GRE)-driven reporter gene assays. HeLa cells were co-transfected with MMTV-Luc, a GRE-driven luciferase, and with a plasmid coding for either isoform 1, isoform 2 or an empty vector as a control (ctr vector). Cells were then treated with increasing concentrations of Dex, and luminescence was measured 48 hours after transfection (Fig. 3b). As expected, GRE activity was enhanced in proportion of Dex concentration in the presence of both isoforms and the control vector (two-way ANOVA Dex effect p < 0.0001). Cells overexpressing isoform 1 showed a significantly lower dose-response curve compared to cells overexpressing either isoform 2 or the control plasmid (two-way ANOVA isoform effect p < 0.0001 and isoform x Dex effect p < 0.0001), which, in turn, were perfectly overlapping. Isoform 1 reduces the activity of GR, meaning that higher concentrations of Dex are required to evoke GR activation. To confirm these findings, the reporter-gene assay was repeated with FKBP51 KO cells. A CRISPR-Cas 9 approach followed by clonal selection was used to generate cells lacking isoform 1 only (iso 1 KO) or both isoforms (full KO) in HeLa cells using a pool of different guide RNAs (see materials section and supplementary Fig. S4). With all genotypes we saw, as expected, a dose-dependent curve in response to Dex (two-way ANOVA Dex effect p < 0.0001). The curve resulting from the luciferase assay in the full KO overlapped with the one from isoform 1-KO (*i.e.* still containing isoform 2). Both KO lines showed an overall increased activity with lower Dex doses as compared to WT (two-way ANOVA isoforms effect p = 0.0008). This result suggests that the lack of isoform 1 increases the sensitivity of GR to Dex, and that isoform 2 alone (as seen in isoform 1 KO) is not able to rescue this effect (Fig. 3c). Taken together, the results of both reporter-gene assays indicate that isoform 1 alone, and not isoform 2, has an inhibitory function on GR.

#### Differential effects on macroautophagy

Next, we proceeded with the analysis of the main macroautophagy markers, since it has been shown that this pathway is regulated by FKBP51 (Gassen *et al*, 2014). Upstream regulation of autophagy is tightly controlled by the kinase AKT. AKT (activated when phosphorylated) inactivates the autophagy initiator BECN1 via phosphorylation. In turn, AKT can be inactivated through dephosphorylation by the phosphatase PHLPP. This latter process is mediated by FKBP51 (Gassen *et al*, 2014). Isoform 1, 2 or an empty vector as control were overexpressed in HeLa cells, and the key markers of macroautophagy were analyzed via western blot. Quantifications of pAKT showed that overexpression of both isoform 1 and 2 led to a decreased phosphorylation of AKT (pAKT) compared to control (Fig. 3d, e). Decreased pAKT leads to an enhanced autophagy, therefore we analyzed the main autophagic markers: BECN1, upstream regulator of autophagy which is modulated directly by AKT, ATG12, involved in the expansion of autophagosomes being covalently bound to ATG5 and targeted to autophagosome vesicles (ATG12-ATG5), and LC3B-II (lipidated form of LC3B-I), marker of autolysosome formation. Overexpression of isoform 1 led to an increase of BECN1 and ATG12-ATG5 (Fig. 3d, f, g). Interestingly, overexpression of isoform 1 did not lead to an increase of LC3BII (normalized on LC3BI) (Fig. 3d, h). Furthermore, overexpression of isoform 2 did not affect levels of BECN1, but led to increased ATG12-ATG5 and LC3BII/I (Fig. 3d, f-h).

#### Differential effects on DNA methyltransferase 1

As we have previously shown, FKBP51 modulates DNA methyltransferase 1 (DNMT1) activity via phosphorylation in response to antidepressants, affecting genome-wide methylation levels (Gassen *et al*, 2015). To test the effect of the two FKBP51 isoforms on the phosphorylation (*i.e.* activation) levels of DNMT1 (pDNMT1), isoforms 1 or 2 of FKBP51 were again overexpressed in HeLa cells. pDNMT1 was detected via western blot analysis and normalized to total DNMT1. Quantifications indicated a large reduction of pDNMT1 in the presence of isoform 1 overexpression (Fig. 3d, i). Contrarily overexpression of isoform 2 did not affect DNMT1 phosphorylation compared to control (Fig. 3d, i).

#### Differential effects on Calcineurin-NFAT signaling

FKBP51 has also been shown to be involved in the regulation of the immune response through Calcineurin-NFAT signaling (Li *et al*, 2002). We analyzed the effect of FKBP51 isoform 1 and 2 overexpression on the phosphorylation of NFAT. Given the importance of a proper immune response, we used the immortalized human T lymphocyte cell line Jurkat for this purpose. Plasmids coding for isoforms 1 or 2 of FKBP51 were overexpressed in Jurkat cells and pNFAT levels were analyzed via western blot. Quantifications revealed an increase of pNFAT when overexpressing isoform 1 (Fig. 3d, j). Conversely, overexpression of isoform 2 did not affect pNFAT levels compared to control (Fig. 3d, j).

Overall, these data revealed that the two FKBP51 isoforms can have equivalent or opposite effects. The reasons behind this and the possible implications will be examined in the discussion part.

## Discussion

With this study we highlighted both commonalities as well as fundamental differences between the two isoforms of the human FKBP51 protein. Using targeted assays, we were able to map differences in Dex responsiveness and half-life of the two isoforms. In fact, in cultured cells as well as in human blood samples, the short variant 4/isoform 2 appears to have a faster turnover, with a more rapid increase in expression upon dexamethasone treatment and a faster protein degradation rate *in vitro*. The faster dynamic that we observe in blood samples compared to HeLa cells, with a return to baseline levels after 23 hours of Dex intake, is most probably due to a metabolization of Dex that occurs *in vivo* but not *in vitro* (Menke *et al*, 2016). The general faster responsiveness of variant 4 mirrors the findings of Marrone and colleagues (Marrone *et al*, 2023a), albeit in a different context, where they observe a similar rapid response of variant 4 when stimulating T cell proliferation as compared to variant 1. Notably, the authors also report a nuclear localization of variant 4, which aligns with our own observations (Fig. 3a). In silico analyses performed with the Expasy Prosite database (https://prosite.expasy.org/) revealed the presence of a putative nuclear localization signal (NLS) inside the unique C-terminal sequence of isoform 2 (supplementary Fig. S1), validating the hypothesis of a selective nuclear localization and function of isoform 2 compared to isoform 1. Collectively, these corroborative findings suggest a potentially unexplored and important role for variant 4 in immediate-response transcriptional processes, warranting further investigation.

On a functional level, our experiments also revealed a partially distinct role for the two isoforms in the different cellular pathways. The two isoforms were found to exert distinct regulatory effects on GR, NFAT and DNMT1 signaling. The regulation of these pathways depends on the interaction of FKBP51 with HSP90. It is therefore not surprising that isoform 1 affects inhibition of GR and phosphorylation of NFAT and DNMT1, while isoform 2 does not have any effect on their function since it lacks the TPR domain responsible for the interaction with HSP90. On the other hand, the autophagic pathway, which is regulated via the interaction of FKBP51 with AKT and PHLPP, is modulated by both isoforms, since the interaction is dependent on the FK1 domain. The immediate consequence of FKBP51‘s interaction to AKT and PHLPP is the dephosphorylation of AKT. Interestingly, while AKT dephosphorylation is equally regulated by the two isoforms (Fig. 3 a), downstream effects, such as increase of autophagy markers, are not (Fig. 3 e, f). This finding suggests the existence of an additional mechanism for which isoform 2 has a decreased effect on autophagy activation. Presumably, isoform 2 has a lower binding affinity for BECN1. Interestingly, though, isoform 2 appears to have a stronger effect in later stages of the autophagic pathway (autophagosome expansion and autolysosome formation), suggesting an alternative pathway, or a faster activity of isoform 2 compared to isoform 1. Once again, these results suggest different functional roles for the two isoforms. Considering the different functions related to the different domains, it would be of particular interest to explore the functions related to the unique C-terminal sequence of 46 aa of isoform 2. While our results show quite distinct functional roles of the two isoforms, the much lower expression of isoform 2 needs to be considered when interpreting overall effects. However, the distinct time dynamic to stimuli, such as activation via glucocorticoids and possibly also other inducers may open time windows in which isoform 2 is present at substantial levels compared to isoform 1. As mentioned above, the functional role of the distinct subcellular location also remains to be explored, as this could also differentially affect biochemical processes in different cellular compartments.

The identification of functional disparities and differences in their dynamic regulation following stimulation between FKBP51 isoforms carries significant weight for future research on FKBP5/51 and for drug design. The understanding that isoform 1 and isoform 2 not only exhibit different responses to dexamethasone but also have distinct roles in regulating the GR suggests that targeted therapies could be developed to modulate these isoforms selectively and the urgency of future studies to address this difference. In terms of future research on FKBP51, these findings emphasize the need for a comprehensive understanding of the roles of different isoforms in various cellular and subcellular contexts and disease states. While animal models have proven invaluable for understanding FKBP5’s role in both physiological and pathological processes, the absence of isoform 2 in rodents may have created a shortsighted gap in our comprehensive understanding of this scaffold protein. Further investigation into the mechanisms underlying the differential functions of isoform 1 and isoform 2, including their distinct interactions with cellular partners and signaling pathways, is essential. This study sheds light on the functional divergence of FKBP51 isoforms and highlights the importance of including such differentiation in future studies. These findings not only contribute to our understanding of FKBP51 biology but also hold promise for the development of isoform-specific, fine-tuned therapeutic strategies in the treatment of a wide range of diseases.

## Materials and Methods

### Reagents and Tools

#### Antibodies

The following primary antibodies were used for western blot: BECN1 (1:1000, Cell Signaling, #3495), ATG12 (1:1000, Cell Signaling, #2010), LC3B-II/I (1:1000, Cell Signaling, #2775), FKBP51 (1:1000, Bethyl, A301-430A and Abcam ab46002), FKBP51 specific for isoform 2 (generously provided by the Maria-Fiammetta Romano lab, Federico II University), AKT (1:1000, Cell Signaling, #4691), pAKT (Ser473 1:1000, Cell Signaling, #4058 and #9275), Actin (1:5000, Santa Cruz Biotechnology, sc-1616), GAPDH (1:8000, Millipore CB1001)

#### Plasmids

FKBP51-FLAG as described in Wochnik et al, 2005).

The following expression vectors were purchased from Promega: FKBP5-pFN21A #FHC02776, GAPDH-pFN21A #FHC02698, HaloTag®-pFN21AB8354 #FHC02776, pFN21A HaloTag® CMV Flexi Vector #9PIG282.

HT-FKBP51 isoform 2 expressing plasmid was generated by enzymatic cloning of the coding sequence (ENST00000542713.1) into the pFN21A HaloTag® CMV Flexi Vector.

FKBP51 CRISPR/Cas9 KO Plasmid (h), sc-401560, consisting of a pool of 3 plasmids, each encoding the Cas9 nuclease and a target-specific 20 nt guide RNA (gRNA) designed for maximum knockout efficiency. Of the 3 plasmids, one contains gRNA targeting exon 11, specific for variant 1, 2 and 3 but not 4, and the other two plasmids contain gRNAs targeting exon 7 and 4, present in all variants.

#### RT-qPCR primers

*FKBP5* variants 1-3 (Exon 11-12), IDT Hs.PT.58.813038: forward primer:

AAAAGGCCAAGGAGCACAAC

reverse primer: TTGAGGAGGGGCCGAGTTC

*FKBP5* all variants (Exon 5-6), IDT Hs.PT.58.20523859 forward primer:

GAACCATTTGTCTTTAGTCTTGGC

reverse primer: CGAGGGAATTTTAGGGAGACTG

*FKBP5* variant 4 (Exon 8-10b), IDT Hs.PT.58.26844122:

forward primer: GAGAAGACCACGACATTCCA

reverse primer: AGCCTGCTCCAATTTTTCTTTG

*YWHAZ* (Exon 9-10), IDT Hs.PT.58.4154200: forward primer:

GTCATACAAAGACAGCACGCTA reverse primer: CCTTCTCCTGCTTCAGCTTC

### Methods and Protocols

#### Cell culture

The HeLa cell line was cultured at 37°C, 6% CO2 in Dulbecco’s Modified Eagle Medium (Gibco) high glucose with GlutaMAX (Thermo Fisher, 31331-028), supplemented with 10% fetal bovine serum (Thermo Fisher, 10270-106) and 1% antibiotic-antimycotic (Thermo Fisher, 15240-062).

The Jurkat cell line was cultured at 37°C, 6% CO2 in RPMI (Gibco) supplemented with 10% FCS and 1% Antibiotic/Antimycotic (Thermo Fisher scientific Inc., Schwerte, Germany)

#### Transfections

Jurkat cells (2 × 10^6^; suspension cells), or with 1x trypsin-EDTA (gibco, 15400-054) detached HeLa cells (2 × 10^6^) were resuspended in 100 μl of transfection buffer [50 mM Hepes (pH 7.3), 90 mM NaCl, 5 mM KCl, and 0.15 mM CaCl_2_]. Up to 2 μg of plasmid DNA was added to the cell suspension, and electroporation was carried out using the Amaxa 2B-Nucleofector system (Lonza). Cells were replated at a density of 105 cells/cm^2^.

For the intracellular localization experiments, Hela cells were transfected with Lipofectamine 3000 transfection reagent (Thermo Fisher, L3000001) according to the supplier’s protocol.

##### Imaging

HeLa cells were seeded on cover cover glasses (Paul Marienfeld, 0117530) and transfected the next day. 24 hours after transfection, cells were live imaged with a Zeiss epifluorescent microscope.

#### Western Blot Analysis

Protein extracts were obtained by lysing cells in 62.5 mM Tris, 2% SDS, and 10% sucrose, supplemented with protease (Sigma, P2714) and phosphatase (Roche, 04906837001) inhibitor cocktails. Samples were sonicated and heated at 95 °C for 5 min. Proteins were separated by SDS-PAGE and electro-transferred onto nitrocellulose membranes. Blots were placed in Tris-buffered saline solution supplemented with 0.05% Tween (Sigma, P2287) (TBS-T) and 5% non-fat milk for 1 hour at room temperature and then incubated with primary antibody (diluted in TBS-T) overnight at 4 °C. Subsequently, blots were washed and probed with the respective horseradish-peroxidase- or fluorophore-conjugated secondary antibody for 1 hour at room temperature. The immuno-reactive bands were visualized either using ECL detection reagent (Millipore, WBKL0500) or directly by excitation of the respective fluorophore. Recording of the band intensities was performed with the ChemiDoc MP system from Bio-Rad.

##### Quantification

All protein data were normalized to Actin or GAPDH, which was detected on the same blot. In the case of AKT phosphorylation, the ratio of pAKTS473 to total AKT was calculated. Similarly, the direct ration of LC3BII over LC3BI is also provided, as well as the ratio over Actin.

#### Real time quantitative polymerase chain reaction (RT-qPCR)

##### HeLa cells

Total RNA was isolated from HeLa cells with the RNeasy mini kit (Qiagen, 74104) following the manufacturer’s protocols. Reverse transcription was performed using SuperScript II reverse transcriptase (Thermo Fisher, 18064014). Subsequently, the cDNA was amplified in triplicates with the LightCycler 480 Instrument II (Roche, Mannheim, Germany) using primers from IDT and TaqManTM Fast Advanced Master Mix (Thermo Fisher, 4444964).

##### Human samples

Total RNA was isolated from whole blood from healthy, all male donors aged 20 to 30 years, administered 1.5 mg dexamethasone per os at 12 pm. Blood draws (Pax-Gene RNA tubes) were repeated right before Dex administration (12 pm) as well as 1, 3, 6 and 23h there-after Details, including IRB approval are described in Wiechmann et al, 2019. Reverse transcription was performed using SuperScript II reverse transcriptase (Thermo Fisher, 18064014). Subsequently, the cDNA was amplified in triplicates with the LightCycler 480 Instrument II (Roche, Mannheim, Germany) using primers from IDT and TaqManTM Fast Advanced Master Mix (Thermo Fisher, 4444964). The 130 samples corresponding to the five time points of the 26 participants were distributed for RT-qPCR to have all the time points for each participant on the same plate and the same assay on the same plate. Samples were run in technical triplicates and standards were sun on each plate to calculate the efficiency. Quality control of the raw data and were efficiency-corrected ΔΔCP method published by Pfaffl (Pfaffl, 2001) were performed with R studio (version 2021.09.1, RStudio Team (2021). RStudio: Integrated Development Environment for R. RStudio, PBC, Boston, MA URL http://www.rstudio.com/). Statistical analyses were performed Prism version 9.0.0 (GraphPad Software, La Jolla California USA, www.graphpad.com)

#### CRISPR-Cas9 KO generation

Generation of *FKBP5* KO HeLa cell line: using Lipofectamine 3000 transfection reagent (Thermo Fisher, L3000001), cells were transfected with a pool of three CRISPR/Cas9 plasmids containing gRNA targeting human *FKBP5* and a GFP reporter (Santa Cruz, sc-401560). 48 hours post transfection, cells were FACS sorted for GFP as single cells into a 96-well plate using BD FACS ARIA III) in FACS medium [PBS, 0.5% BSA Fraction V, 2 mM EDTA, 20mM Glucose, and 100 U/mL Penicillin-Streptomycin]. Single clones were expanded and western blotting was used to validate the knockouts and variants-specific knockouts were selected based on western blot analyses using antibodies specific for Isoform 1 or 2 as detailed above.

#### Reporter gene assays

For the MMTV-luc reporter gene assay, cells were seeded in 96 well plates in medium containing 10% charcoal-stripped, steroid-free serum and cultured for 24 h before transfection using Lipofectamine 2000 as described by the manufacturer. Unless indicated otherwise, the amounts of transfected plasmids per well were 60 ng of steroid responsive luciferase reporter plasmid MMTV-Luc, 5–7.5 ng of Gaussia-KDEL expression vector as control plasmid, and up to 300 ng of plasmids expressing FKBP51-HTv1; FKBP51-HTv4. If needed, empty expression vector was added to the reaction to equal the total amount of plasmid in all transfections. 24 h after transfection, cells were cultured in fresh medium supplemented with Dex as indicated or DMSO as control for 24 h. To measure reporter gene activity cells were washed once with PBS and lysed in 50 µl passive lysis buffer (0.2% Triton X-100, 100 mM K2HPO4/KH2PO4 pH 7.8). Firefly and Gaussia luciferase activities were measured in the same aliquot using an automatic luminometer equipped with an injector device (Tristar, Berthold). Firefly activity was measured first by adding 50 µl Firefly substrate solution (3 mM MgCl2, 2.4 mM ATP, 120 µM D-Luciferin) to 10 µl lysate in black microtiter plates. By adding 50 µl Gaussia substrate solution (1.1 M NaCl, 2.2 mM Na2EDTA, 0.22 M K2H PO4/KH2PO4, pH 5.1, 0.44 mg/ml BSA, Coelenterazine 3 µg/ml) the firefly reaction was quenched and Gaussia luminescence was measured after a 5 s delay. Firefly activity data represent the ratio of background corrected Firefly to Gaussia luminescence values. The fold stimulation reached at saturating concentrations of hormone was about 3 nM, which is in the range of previous publications (Touma *et al*, 2011; Schülke *et al*, 2010).

#### Pulse chase assay

48 hours after transfection with HaloTag®-tagged plasmids, cells were labeled with HT fluorescent ligands (HaloTag® R110Direct Ligand, Promega) for 24 hours after which the fluorescent ligand was washed off (chase) for the indicated amounts of time. Cells were harvested, proteins extracted minimizing light exposure and western blots were performed. Fluorescence was successively measured on membrane with the ChemiDoc MP system from Bio-Rad.

### Ethics approval and consent to participate

All studies with human samples were approved by the local ethics committee of the Medical School of the Ludwig Maximilians University, and all participants gave informed consent.

